# Generally applicable transcriptome-wide analysis of translational efficiency using anota2seq

**DOI:** 10.1101/106922

**Authors:** Christian Oertlin, Julie Lorent, Valentina Gandin, Carl Murie, Laia Masvidal, Marie Cargnello, Luc Furic, Ivan Topisirovic, Ola Larsson

## Abstract

mRNA translation plays an evolutionarily conserved role in homeostasis and when dysregulated results in various disorders. Optimal and universally applicable analytical methods to study transcriptome-wide changes in translational efficiency are therefore critical for understanding the complex role of translation regulation under physiological and pathological conditions. Techniques used to interrogate translatomes, including polysome- and ribosome-profiling, require adjustment for changes in total mRNA levels to capture *bona fide* alterations in translational efficiency. Herein, we present the anota2seq algorithm for such analysis using data from ribosome- or polysome-profiling quantified by DNA-microarrays or RNA sequencing, which outperforms current methods for identification of changes in translational efficiency. In contrast to available analytical methods, anota2seq also allows capture of an underappreciated mode for regulation of gene expression whereby translation acts as a buffering mechanism which maintains constant protein levels despite fluctuations in mRNA levels (“translational buffering”). Application of anota2seq shows that insulin affects gene expression at multiple levels, in a largely mTOR-dependent manner. Moreover, insulin induces levels of a subset of mRNAs independently of mTOR that undergo translational buffering upon mTOR inhibition. Thus, the universal anota2seq algorithm allows efficient and hitherto unprecedented interrogation of translatomes and enables studies of translational buffering which represents an unexplored mechanism for regulating of gene expression.

## INTRODUCTION

Regulation of gene expression is a multi-step process including transcription, mRNA-processing, -transport, -stability, -translation and protein stability ^1^. Although the precise relative contribution of each of these processes to observed gene expression remains controversial ^2^ and context dependent ^3,4^, several studies have implicated mRNA translation as a key mechanism that determines the composition of the proteome ^5,6^. Notably, rapid adaptation to changes in the cellular environment requires precipitous adjustment of the proteome which is, in addition to protein degradation, largely accommodated by altering mRNA translation ^7^. Direct transcriptome-wide quantification of mRNA translation is therefore required to enhance the understanding of how protein levels are regulated in response to a variety of stimuli and stressors and in normal vs. diseased cells.

Translation is thought to be predominantly regulated at the level of initiation ^8^. Translational efficiency is therefore reflected by the proportion of total mRNA that is associated with efficiently translating ribosomes (polyribosomes or polysomes). Thus, the number of ribosomes associated with mRNA is proportional to translation efficiency ^9^ and generally parallels corresponding protein levels (**Fig. 1A**). Polysome-profiling separates mRNAs based on the number of ribosomes that they are associated with ^10^. Such polysome-associated and total mRNA are then quantified in parallel using DNA-microarrays or RNA sequencing (RNAseq) ^7^. During polysome profiling, efficiently translated mRNAs are commonly defined as those associated with >3 ribosomes, (hereafter called polysome-associated mRNA). A potential pitfall with setting a >3 ribosomes threshold for efficiently translated mRNA is that shifts in ribosome-association that occur exclusively in fractions containing >3 (or <3) ribosomes, although possibly biologically meaningful, will not be captured. Empirical evidence, however, justifies this approach as upon a change in translational efficiency, the normally distributed population of mRNA copies from a single gene at least partly shifts across the >3 ribosome threshold under a variety of experimental conditions ^11^. Indeed, alterations in translational efficiency can be broadly divided in two categories: those that occur when mRNAs shift from polysomes to sub-polysomal fractions (i.e. “on-off” regulation) or those where the shift in ribosome-association occur within the polysomes ^11^. The former scenario is illustrated by mTOR-dependent regulation of 5’-terminal oligopyrimidine motif (TOP) mRNA translation. TOP mRNAs encode components of the translational machinery and are amongst subset of transcripts which are associated with the heaviest polysomes under conditions wherein the mammalian/mechanistic target of rapamycin (mTOR) is active ^11^. Inhibition of mTOR results in their near complete dissociation from ribosomes ^11^. In stark contrast, under conditions when mTOR is activated, mRNAs encoding proliferation-promoting proteins (e.g. cyclins) are associated with fewer ribosomes relative to TOP mRNAs, and upon mTOR inhibition the reduction in their ribosome association is largely contained within polysomes and is hence less dramatic as compared to TOP mRNAs ^11^. Importantly, both modes of translational regulation result in shifts across the >3 ribosome threshold and thus lead to changes in amount of mRNA associated with >3 ribosomes. Of note, analysing mRNA content in each polysomal fraction (up to 10 fractions are typically collected) is associated with unrealistic cost and time burden ^12^.

**Figure 1.**
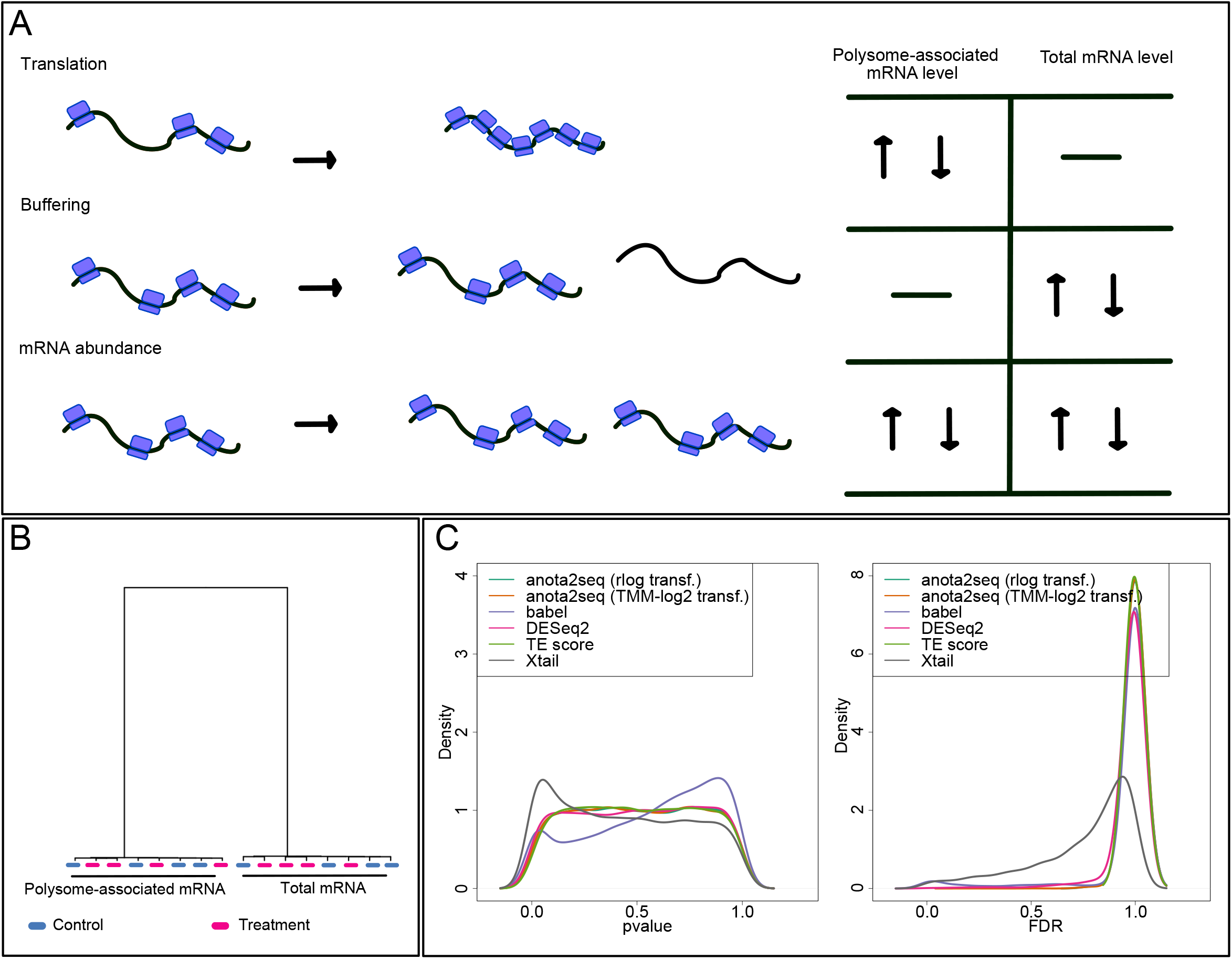
Analysis of differential translation using Xtail or babel is associated with increased false positive findings. (**A**). Gene expression can be modulated via changes in translational efficiency leading to altered protein levels or buffering where the number of mRNA copies for a single gene changes but the number of such copies that are efficiently translated is unaltered; and via changes in mRNA abundance paralleled by changes in levels of efficiently translated mRNA. Statistical methods that distinguish between changes in translational efficiency that affect protein levels from those that do not are therefore warranted. (**B**) Hierarchical clustering of a simulated treatment and control dataset including polysome-associated mRNA and total mRNA levels where no mRNAs are regulated (i.e. a NULL model). (**C**) P-value and FDR density plots for analysis of differential comparing treatment to control for each method using the data from (B).

Ribosome-profiling involves isolation of ribosome-protected fragments (RPFs), generated by RNase treatment, irrespective of the number of ribosomes bound to each individual mRNA molecule ^13^. Ribosome-profiling thereby, in conjunction with RNAseq, maps positions of ribosomes on mRNAs with a single nucleotide resolution ^13–15^. As a result, ribosome-, but not polysome-profiling provides information on regions of an mRNA that are associated with ribosomes. For example, ribosome-but not polysome-profiling allows discrimination between translation of an upstream open reading frame (uORF) and a major ORF ^16^. Indeed, SLC45A4 shows a shift in translation of the uORF to the ORF upon arsenite treatment, a regulatory event captured by ribosome-profiling, which cannot be resolved using polysome-profiling ^16^. It has recently, however, been reported that interpretation of ribosome profiling data may be challenging due to biases introduced by commonly used translation inhibitors including cycloheximide, the RNAse used to generate RPFs and/or lack of appropriate quality control ^17–21^. Application of ribosome-profiling also leads to greater biases as compared to polysome-profiling when employed to analyze changes in translational efficiency, which is a consequence of a strong dependence of ribosome profiling on the magnitude of the shift of the mRNA across the polysomes, as well as mRNA abundance ^11,22^. This dramatically favors identification of highly abundant mRNAs showing large shifts in translational efficiency (e.g. TOP mRNAs) over those of lower abundance that exhibit lesser shifts (e.g. cyclins) when applying ribosome-profiling ^11,22^. Thus, while ribosome-profiling provides information on ribosome positioning on mRNA, and was critical for understanding a plethora of underexplored aspects of protein synthesis ^14,15^, polysome-profiling appears to be less biased by the magnitude of shifts in translational efficiency and mRNA abundance when analyzing translational efficiency on a transcriptome-wide scale ^11,22^. We will therefore discuss transcriptome-wide analysis of translational efficiency using data obtained from the polysome-profiling strategy, but notably anota2seq can be equally applied to ribosome-profiling data.

Amounts of polysome-associated mRNAs are also influenced by steps in the gene expression pathway that act upstream of translation which modulate total mRNA levels (e.g. transcription and mRNA stability; **Fig. 1A**). Analysis of *bona fide* changes in translational efficiency therefore seeks to identify changes in the amount of polysome-associated mRNAs that are independent of changes in total mRNA levels. To this end the <u>an</u>alysis <u>o</u>f <u>t</u>ranslational <u>a</u>ctivity (anota) algorithm ^23^ applies per-gene analysis of partial variance (APV) ^24^ coupled with variance shrinkage ^25^. This approach is superior to methods comparing differences between log ratios (i.e. between polysome-associated mRNA and total mRNA, commonly referred to as translational efficiency scores [TE]) inasmuch as log ratio based approaches do not efficiently adjust changes in polysome-associated mRNA for changes in total mRNA levels due to a phenomenon referred to as spurious correlation ^26^. Spurious correlations were initially described by K. Pearson and imply that a difference score, which in this case is the log ratio of polysome-associated mRNA (or RPF in the case of ribosome-profiling) to total mRNA levels, can systematically correlate with total mRNA levels ^27^. A change in total mRNA levels may therefore lead to false positive identification of differential translation and biological misinterpretations. Unfortunately, such spurious correlations are abundant in both polysome- and ribosome-profiling studies suggesting that log ratio based approaches should be avoided ^26^. A recent study ^28^ evaluated anota on data generated using RNAseq and implied poor performance for such data. Anota however, was developed for normalized data on a continuous logarithmic scale, and thus cannot be applied on non-transformed count data originating from RNAseq studies. This highlights that inappropriate use of analytical methods is an emerging issue in quantitative biology, and in particular in systems biology approaches.

Several methods have been specifically developed for analysis of differential translation using RNAseq data as input and are available within the commonly used statistical programming language “R”, including babel ^29^ and Xtail ^28^; or were developed for analysis of differential expression using RNAseq data and can be utilized for analysis of differential translation, including DESeq2 ^30^. Because all these methods use log-ratios (or equivalent analysis) and therefore are likely to entail spurious correlations, we aimed to develop an anota-like approach for count data (i.e. RNAseq). Initially we considered an approach based on edgeR ^31^ or DESeq2 ^30^. However, because these methods cannot apply gene specific covariates, this was unachievable ^30,31^. Instead, we identified suitable data normalization/transformation for RNAseq data allowing application of APV (i.e. passing quality control steps) ^23^. Essentially, methods for analysis of changes in translational efficiency are distinct from those that aim to identify translated regions using ribosome profiling data as input including RiboTaper ^32^, RibORF ^33^ and ORFscore ^34^.

We studied effects of insulin on gene expression in MCF7 cells using polysome-profiling quantified by DNA-microarrays or RNAseq in parallel. This allowed us to simulate a realistic data set to compare the performance of the various methods for analysis of changes in translational efficiency, which indicated superior performance of anota2seq. These data were also employed to establish key quantification- and normalization/transformation-related aspects that allow broad application of anota2seq, and directly compared the performance of DNA-microarrays to RNAseq. This suggested that RNAseq allow for identification of changes in translational efficiency across a broader expression range than DNA-microarrays. Moreover, using anota2seq, we show that in addition to changes in translational efficiency that modulate protein levels, translational buffering, which maintains protein levels constant under conditions wherein mRNA levels are altered, is a pervasive mode of regulation of gene expression in mammals. Translational buffering has previously been suggested to maintain protein levels constant despite inter-individual variation in mRNA levels between samples from different patients ^35^ and different yeast strains ^36–38^. Herein, we show that a many mRNAs are translationally buffered during an acute response to a physiological stimulus (insulin) and inhibition of a key signalling pathway (mTOR). Notably, analytical methods other than anota2seq failed to discriminate between changes in translational efficiency leading to altered protein levels and buffering.

## RESULTS

### Babel and Xtail algorithms underperform under a NULL model

To evaluate the performance of methods for analysis of translatomes quantified by RNAseq, we employed a simulated data approach (see materials and methods for details; **Fig. S1**). By monitoring quality control steps (see materials and methods), we first identified suitable data normalization/transformation for application of anota2seq (rlog ^30^ and TMM-log2 ^39^). We then compared the performance of anota2seq using rlog or TMM-log2 transformed data to babel ^29^, DESeq2 ^30^, translational efficiency (TE) score and Xtail ^28^. An essential aspect during identification of differential expression is the control of type I error/false discovery rates (FDR) under a NULL model (i.e. when there are no true differences in gene expression). Notably, the performance under the NULL model is unrelated to the sensitivity of the method. Under the NULL model the distribution of p-values should be uniform resulting in FDRs for all mRNAs to equal 1 (i.e. after adjusting for multiple testing). We therefore simulated a dataset with two conditions sampled from the same distribution (i.e. where there were no differences in expression between conditions). Consistent with observations from empirical data sets ^40^, simulated data for total mRNA or polysome-associated mRNA differed when assessed using hierarchical clustering (**Fig. 1B**). Moreover, as expected under a NULL model, there was no separation of the treatment and control groups (**Fig. 1B**). The resulting density plots of p-values for changes in translation revealed uniform distributions for all methods except Xtail, which has an overrepresentation of low p-values, and babel, which shows an overrepresentation of high p-values and a local enrichment of low p-values under the NULL model (**Fig. 1C**). Accordingly, both Xtail and babel assign low FDRs to a subset of mRNAs even when analysing data sets which were *a priori* set to be devoid of any changes in translational efficiency (**Fig. 1C**). Xtail and babel therefore have limited usability as they indicate changes in translational efficiency even when such changes are absent in the simulated data set.

### Anota2seq outperforms current methods for identification of changes in translational efficiency affecting protein levels

Changes in levels of polysome-associated mRNAs are thought to follow two major patterns: i) changes in total mRNA abundance caused by modulation of upstream steps in the gene expression pathway (e.g. transcription and mRNA stability) that results in congruent alterations in the levels of polysome-associated mRNAs (**Fig. 1A**) and ii) variations in levels of polysome-associated mRNA in the absence of fluctuations in total mRNA levels, which represent changes in translation efficiency (**Fig. 1A**). Both patterns of regulation result in changes in protein levels. More recently, it has been proposed that translation can “buffer” changes in mRNA levels to maintain constant levels of proteins ^17,36,37,41^. During translational buffering, alterations in total mRNA levels are not accompanied by changes in their polysome-association, and thus although mRNA levels change, the levels of corresponding proteins remain constant (**Fig. 1A**). To evaluate the performance of the various methods, we simulated a data set under two conditions corresponding to control and treatment with all three regulatory patterns (537 mRNAs per pattern along with 9137 unchanged mRNAs): changes in mRNA abundance; and translational efficiency resulting in altered protein levels or buffering (**Fig. 2A**). Hierarchical clustering showed the expected pattern whereby polysome-associated mRNA and total mRNA cluster separately. Moreover, treatment and control groups form clusters within each of these RNA-source related clusters. The simulated data set therefore captures the complex structure of polysome-profiling data (which is similar to ribosome-profiling data ^40^; **Fig. 2B**). We next determined the performance of each method in detecting changes in translational efficiency affecting protein levels. Accordingly, identification of mRNAs belonging to the translation group (**Fig. 1A**) was considered true positive events, whereas identification of unchanged mRNAs; and mRNAs from the buffered (Fig. 1A) and mRNA abundance (Fig. 1A) groups were considered as false positive events (Fig. 2A). To assess the performance of the methods, we used area under the curve (AUC) and partial AUC (pAUC) of receiver operator curves (ROC) ^42^. The methods were applied using default settings on 5 replicate data sets simulated as in **Fig. 2A** (to assess the variability of the simulation) followed by assessment of their outputs prior to any filtering (see material and methods). ROC curves show that anota2seq analysis on rlog or TMM-log2 data performs equally well and outperforms all other methods as judged by AUC and pAUCs (**Fig. 2C**, table 1). In addition, precision recall curves reveal high initial precision values for anota2seq compared to other methods (**Fig. 2C**). This can be explained by the analysis principle of the other methods. TE-score, babel, DESeq2 and Xtail cannot separate changes in translational efficiency affecting protein levels (**Fig. 1A**) from buffering (**Fig. 1A**) whereas this distinction is made using anota2seq. This is consistent with the reported superior performance of anota as compared to TE-score in reflecting changes in the proteome ^43^.

**Figure 2.**
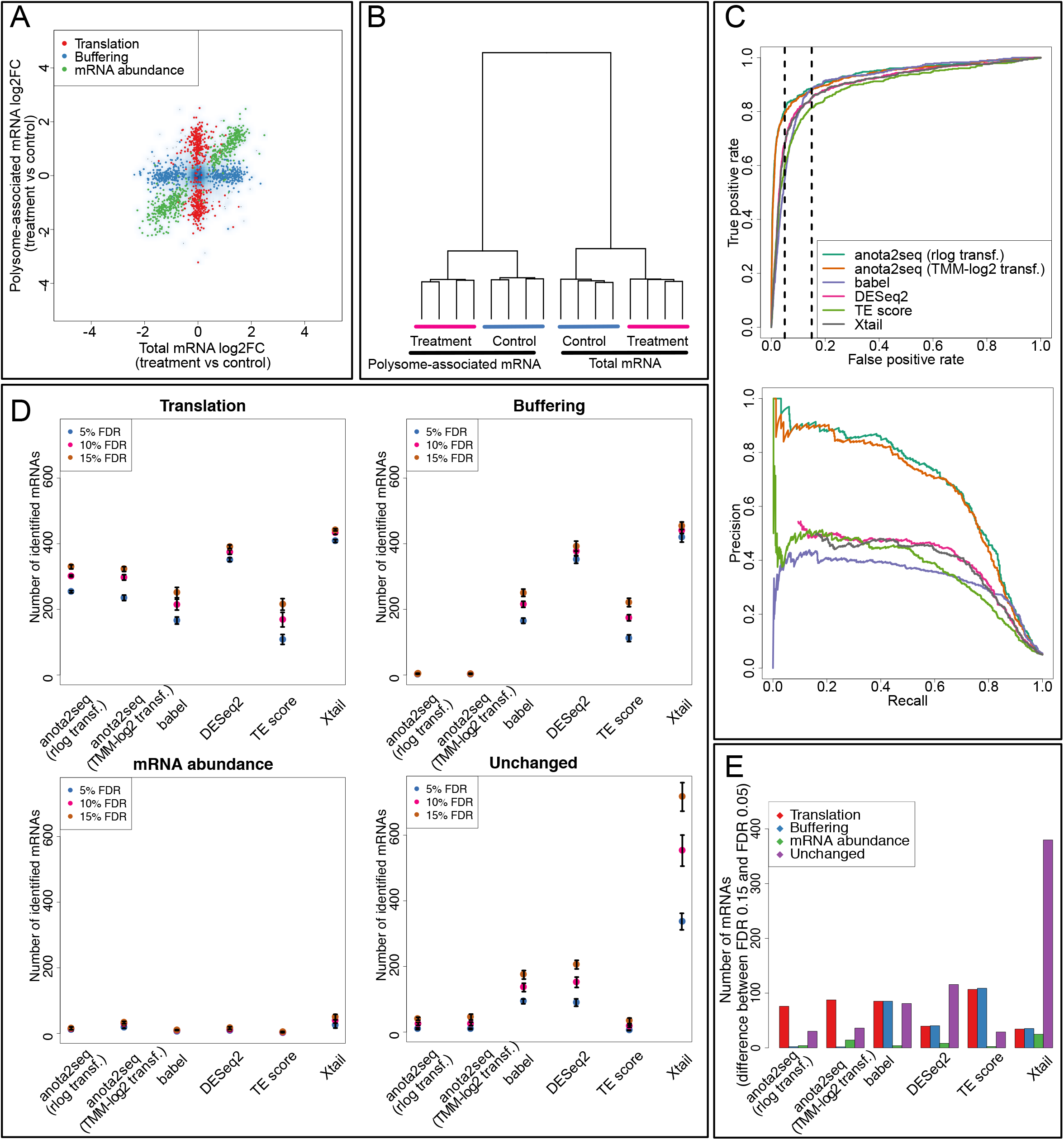
Anota2seq outperforms other methods for analysis of changes in translational efficiency affecting protein levels. (**A**) Scatterplot of polysome-associated and total mRNA log2 fold changes between treatment and control groups for a simulated data set. Simulated mRNAs regulated by changes in mRNA abundance; and translational efficiency affecting protein levels or buffering are indicated. (**B**) Hierarchical clustering of gene expression data from (A). (**C**) Receiver operator curves for analysis of differential translation in a simulated dataset (i.e. from A-B; top). Precision recall curves for analysis of differential translation in the simulated dataset (bottom). For both analyses, identification of a gene simulated as changing its translation leading to altered protein levels (Fig. 1A) was considered a true positive event. Vertical lines indicate 5% and 15% false positive rates. (**D**) Numbers of mRNAs identified as differentially translated that belonged to the simulated groups with changes in translational efficiency leading to altered protein levels (translation, true positives [TP]) or buffering (false positives [FP]); changes in mRNA abundance (FP) or were unchanged (FP) are indicated for each method at several FDR thresholds (mean and standard deviations from 5 simulated data sets are indicated). (**E**) Differences in the number of mRNAs showing differential translation belonging to the 4 categories in (D) when changing the FDR threshold from 5% to 15% are shown (mean from 5 simulated data sets).

**Table 1.** Mean and standard deviation (sd) of pAUCs and AUCs for an ROC analysis assessing performance for identification of changes in translational efficiency affecting protein levels in simulated datasets (n=5) including unchanged mRNAs and groups of mRNAs with changes in translational efficiency affecting protein levels or buffering; or mRNA abundance.

| Method | n | pAUC FPR 5% |  | pAUC FPR 15% |  | AUC |  |
| --- | --- | --- | --- | --- | --- | --- | --- |
|  |  | mean | sd | mean | sd | mean | sd |
| anota2seq (rlog transf.) | 5 | 0.6700 | 0.0159 | 0.8073 | 0.0140 | 0.9486 | 0.0069 |
| anota2seq (TMM-log2 transf.) | 5 | 0.6557 | 0.0178 | 0.7972 | 0.0166 | 0.9444 | 0.0069 |
| babel | 5 | 0.2860 | 0.0144 | 0.5930 | 0.0147 | 0.9124 | 0.0044 |
| DESeq2 | 5 | 0.4080 | 0.0106 | 0.6679 | 0.0052 | 0.9145 | 0.0053 |
| TE score | 5 | 0.3602 | 0.0201 | 0.6072 | 0.0105 | 0.8945 | 0.0050 |
| Xtail | 5 | 0.3893 | 0.0117 | 0.6563 | 0.0056 | 0.9136 | 0.0059 |

To further characterize the performance of the methods at commonly employed FDR thresholds we counted the number of identified mRNAs from each regulatory group (translation, buffering and mRNA abundance; **Fig. 1A**) and unchanged mRNAs at a 5%, 10% or 15% FDR threshold (**Fig. 2D**; using the default settings of each method). Babel and TE score identified fewer true positive events (i.e. changes in translational efficiency affecting protein levels) than the other methods. Anota2seq identified approximately the same number of true positives at the 15% FDR threshold as DESeq2 while Xtail identified slightly more true positive events. The number of mRNAs identified from the set of buffered mRNAs (i.e. false positives) by each method reveals that only anota2seq can efficiently distinguish between changes in translational efficiency, which is expected to affect protein levels, from buffering, which maintains constant protein levels (**Fig. 2D**). Notwithstanding that all methods perform similarly in terms of rejecting mRNAs that show changes in mRNA abundance (i.e. congruent changes in total and polysome-associated mRNA; i.e. false positives), there were dramatic differences in terms of identification of unchanged mRNAs (i.e. false positives). Anota2seq and TE-score identified few such mRNAs while babel and DEseq2 identified a sizeable number. Strikingly, Xtail identified >600 such mRNAs at FDR<15%. The latter finding is consistent with the poor performance of Xtail under the NULL model (**Fig. 1C**). To contrast a 5% and 15% FDR threshold, we calculated the difference in number of identified mRNAs from each regulatory pattern (**Fig. 2E**). For anota2seq, there was a gain in true positives at the cost of a smaller increase in false positives, whereas for other methods, especially Xtail, increasing the FDR threshold introduces more false positives than additional true positives.

### Anota2seq outperforms current methods for identification of changes in translational efficiency affecting protein levels also in the absence of translational buffering

The above simulation included equal number of mRNAs regulated via changes in mRNA abundance, translational efficiency affecting protein levels and buffering (**Fig. 1A, 2A**). Depending on the condition, however, changes in translational efficiency affecting protein levels or buffering may predominate and hence it was of interest to compare the methods using a simulated data set containing changes in translational efficiency affecting protein levels (537 mRNAs) and mRNA abundance (537 mRNAs) groups along with unchanged mRNAs (9674 mRNAs), but without the buffering group (**Fig. S2A-B**). As expected from their inability to separate translation and buffering groups (**Fig. 2D**), under these conditions babel, DESeq2, TE score and Xtail showed improved performance as judged by pAUC and AUC, which was comparable to anota2seq (**table S1**). Moreover, removal of the buffering group resulted in an increase in the precision (compare **Fig. S2C** to **Fig 2C**). As expected, the performance under different FDR thresholds paralleled the full simulation (i.e. also including the buffering group; compare **Fig. S2D-E** to **Fig. 2D-E**). This included comparable performance of anota2seq and DEseq2 ^30^ for identification of mRNAs from the translation group (**Fig. 1A**) and elevated identification of unchanged mRNAs (i.e. false positives) for babel, DESeq2 and Xtail. Analogous to the full simulation, TE-score identified fewer mRNAs from the translation group but did not assign unchanged mRNAs low FDRs. These results demonstrate that while anota2seq shows superior performance under conditions wherein gene expression is regulated via changes in translational efficiency affecting protein levels and buffering (**Fig. 2D**), it also outperforms other methods in the absence of buffering (**Fig. S2D**). Therefore, anota2seq can be applied to efficiently identify changes in translation, which correspond to alterations in protein levels, independent of what the underlying regulatory patterns are.

These results appear to contradict a recent report which suggested good ROC and precision/recall performance for babel, TE score and Xtail and poor performance for anota ^28^. While the reported poor performance of anota ^28^ was caused by inappropriate application of anota on non-transformed counts, the difference in precision recall performance for babel, TE score and Xtail was unclear. We therefore examined the simulated data set used during development of Xtail in detail (**Fig. S3A**). This revealed that mRNAs selected as true positives for changing their translational efficiency belonged to any regulated group (translation, buffering or mRNA abundance [**Fig. 1A**]) and also included mRNAs with seemingly unchanged expression (**Fig. S3B** top left). Moreover, there were mRNAs which showed increased polysome-association but strongly decreased mRNA levels, which represent unlikely biological events that were not observed in any of the empirical data sets examined ^28,44^ (**Fig. S3B** top right and lower panels; notably this is distinct from buffering wherein levels of polysome-associated mRNA remain largely unchanged despite alterations in mRNA levels). We therefore reclassified mRNAs (**Fig. S3C**). We next examined the population of mRNAs that were identified at different FDR thresholds with largely similar results as observed for our simulated datasets, except for babel, which identified very few regulated events (**Fig. S3D**). Thus, the optimal performance in analysis of changes in translation leading to altered protein levels observed for anota2seq is not limited to the herein simulated dataset.

### Analysis of translational buffering using anota2seq

In addition to changes in translational efficiency leading to altered protein levels, buffering also holds potentially important information about translational control as it is thought to function as a mechanism that maintains protein levels despite changes in mRNA expression ^35,37^. Methods to capture buffering, however, have not been developed to date and we therefore implemented such analysis in the anota2seq software. This implementation is based on the same principle as anota2seq analysis that captures changes in translational efficiency affecting protein levels (i.e. APV coupled with variance shrinkage) except that it captures changes in total mRNA levels that are not paralleled by changes in levels of polysome-associated mRNA. To assess the performance of anota2seq for analysis of buffering, we used the same data set as in **Fig. 2A-B** (i.e. with equal numbers of mRNAs from the translation, buffering and mRNA abundance groups; **Fig 1A**). In contrast to the analysis in figure **2A-B**, identification of mRNAs from the buffering group was considered as true positive events, whereas identification of unchanged mRNAs or mRNAs belonging to translation and mRNA abundance groups were considered false positive events. As expected, pAUC and AUC for analysis of buffering were comparable to the performance of anota2seq for analysis of translational efficiency affecting protein levels (**Fig. 3A** and **table 2**). Critically, very few mRNAs belonging to the translation group were identified during analysis of buffering (**Fig. 3B**). Moreover, a relaxed FDR threshold primarily led to additional identification of mRNAs from the buffered group (**Fig. 3C**). Thus, anota2seq can be efficiently applied to identify mRNAs that are buffered at the level of translation.

**Figure 3.**
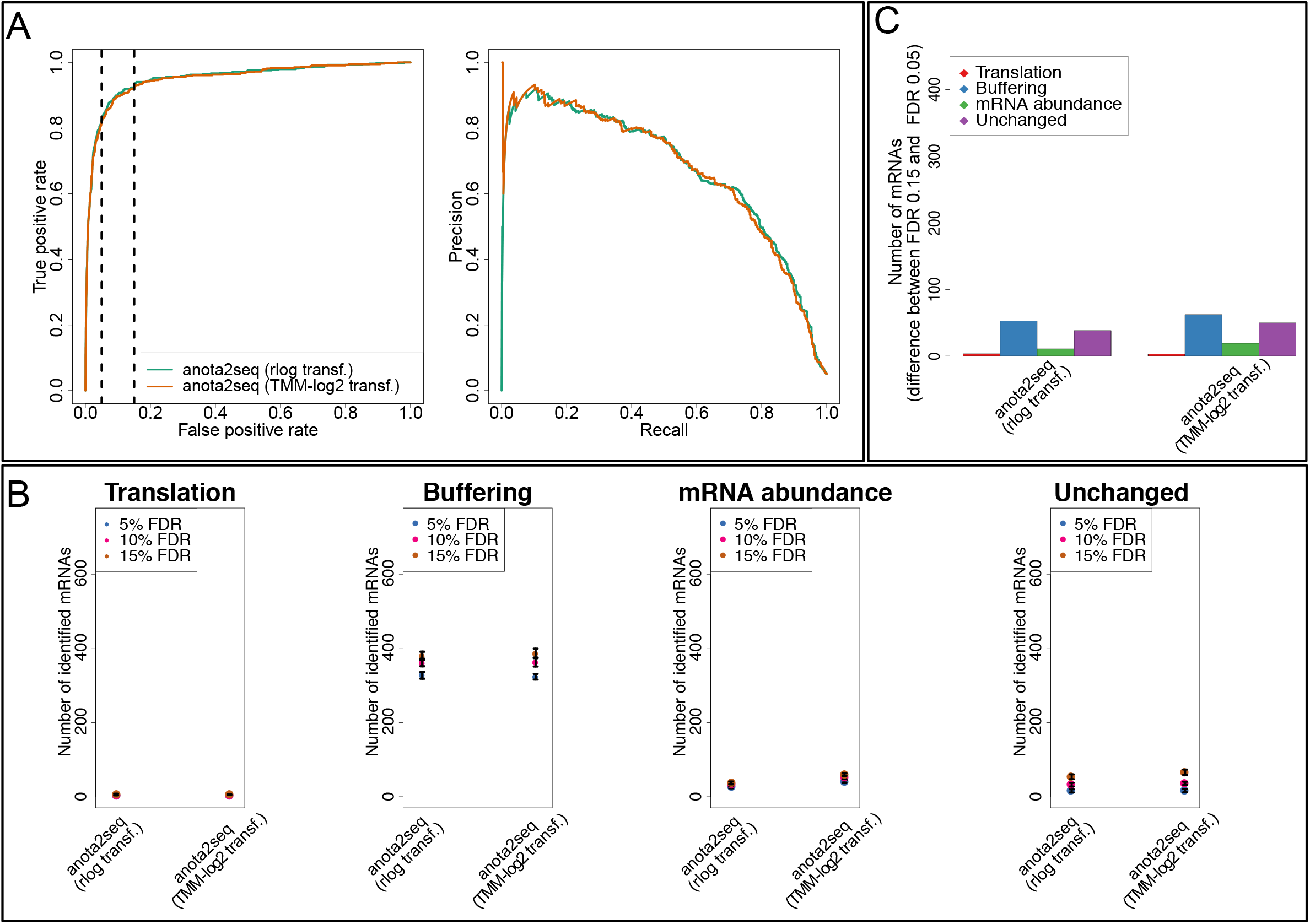
Efficient identification of translational buffering using anota2seq. (**A**) Receiver operator curve (left) and precision recall curve (right) for analysis of translational buffering using anota2seq on a simulated dataset. The dataset is the same as in Fig. 2A-B but identification of mRNAs from the simulated buffering group were considered true positive events. Vertical lines indicate 5% and 15% false positive rates. (**B**) Numbers of mRNAs identified as buffered belonged to the simulated groups with changes in translational efficiency leading to altered protein levels (FP) or buffering (TP), changes in mRNA abundance (FP) or were unchanged (FP) are indicated for each method at several FDR thresholds (mean and standard deviations from 5 simulated data sets are indicated). (**C**) Difference in the number of mRNAs identified as buffered belonging to the 4 categories in (B) when changing the FDR threshold from 5% to 15% (mean from 5 simulated data sets).

**Table 2.** Mean and standard deviation (sd) of AUCs and pAUCs for an ROC analysis assessing performance for identification of changes in translational efficiency leading to buffering in simulated datasets (n=5) including unchanged mRNAs and groups of mRNAs with changes in translational efficiency affecting protein levels or buffering; or mRNA abundance.

| Method | n | pAUC FPR 5% |  | pAUC FPR 15% |  | AUC |  |
| --- | --- | --- | --- | --- | --- | --- | --- |
|  |  | mean | sd | mean | sd | mean | sd |
| anota2seq (rlog transf.) | 5 | 0.651 | 0.021 | 0.805 | 0.021 | 0.952 | 0.010 |
| anota2seq (TMM-log2 transf.) | 5 | 0.646 | 0.021 | 0.800 | 0.021 | 0.949 | 0.011 |

### Assessing robustness against noise and varying sequencing depth

Experimental and technical challenges and/or study setups may give rise to datasets exhibiting dramatically different characteristics. This includes different noise level and varying sequencing depth ^45,46^. To test the effects of variations in noise or sequencing depth on performance of translatome analyses, we simulated datasets with increasing noise levels (5, 10 or 15%; **Fig. S4A**) or increasing number of RNAseq reads (5, 10, 15 or 25 million reads mapping to mRNAs per sample). Increased noise lead to a moderate decrease in the number of true positives identified at an FDR of 15% for all methods except TE score based analysis, which was particularly unstable (**Fig. S4B-C**). This is associated with a decrease in performance as assessed by ROC and precision recall curves (**Fig. S5A-F**). Judged by pAUC and AUC, anota2seq outperforms other methods at all noise levels in analysis of changes in translational efficiency affecting protein levels using simulated data sets including all regulatory groups (i.e. translation, buffering and mRNA abundance [**Fig 1A**]; **Fig. S5; table S2**). Reduced sequencing depth had no clear effect on ROC analysis within methods (**Fig. S6**). Nevertheless, the number of identified true positive events decreased by ~10–50 mRNAs with reduced sequencing depth (**Fig. S7A-B**). In our data set about 49% of the sequenced reads mapped uniquely to genome regions of mRNAs coding for proteins (**Fig. S9A**). As 5 million such reads appears sufficient for analysis, a low but adequate sequencing depth is >10 million reads although this is likely to depend on sample quality and protocol used for preparation of RNAseq libraries.

### Incorporating batch adjustment in anota2seq

Hybridization based DNA-microarrays and count based RNAseq are commonly used to quantify mRNA levels. Both platforms have their intrinsic strengths and weaknesses. For example, DNA microarrays have higher background signals but are less susceptible to issues associated with very high expression of only a limited number of mRNAs that may dominate the output, as sometimes seen in RNAseq ^47^. Nevertheless, it remains to be determined how these technical traits affect transcriptome-wide analysis of the translatome.

To directly test the performance of anota2seq on datasets obtained via DNA-microarrays and RNAseq, we performed a polysome-profiling study focusing on the effects of insulin on mRNA abundance; and translational efficiency affecting protein levels and buffering. We also assessed the dependence of such changes in gene expression on mTOR complex 1 (mTORC1). mTORC1 is activated by insulin via the PI3K/AKT pathway and acts as a major anabolic pathway in the cell that bolsters lipid, protein and nucleotide synthesis ^48–50^. To this end, MCF7 cells were serum starved for 16 hours and treated with a vehicle (DMSO; control), insulin (4.2nM) or insulin (4.2nM) in combination with the active-site mTOR inhibitor torin1 (250nM) for 4 hours (**Fig. 4A**). Despite harbouring an activating PIK3CA mutation ^51^, mTORC1 signalling in MCF7 cells is highly responsive to insulin as illustrated by a dramatic induction in phosphorylation of mTORC1 downstream targets eIF4E-binding protein 1 (4E-BP1) together with ribosomal protein S6 kinases (S6K) and its downstream target ribosomal protein S6 (rpS6) which were abolished by the active site mTOR inhibitor torin1 (**Fig. S8A**). Absorbance profiles revealed that, as expected, insulin induced mRNA translation globally, as evidenced by a decrease in the monosome peak and concomitant increase in polysome area as compared to control, which was reversed by torin1 (**Fig. S8B**). Total and polysome-associated mRNA (>3 ribosomes) were isolated and quantified in parallel using DNA-microarrays and RNAseq to generate two data sets from the exact same samples.

**Figure 4.**
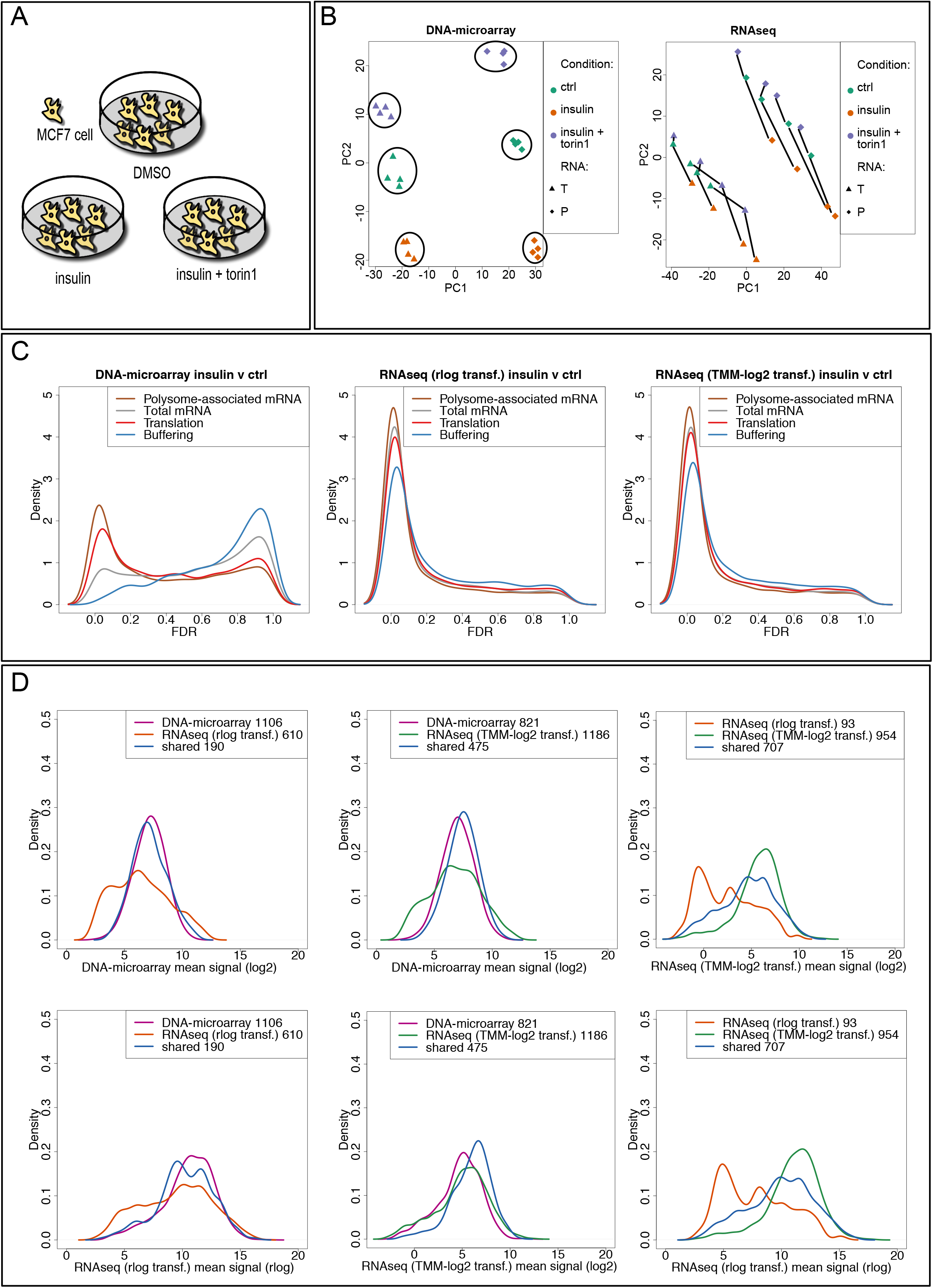
Platform and data normalization/transformation characteristics during anota2seq analysis. (**A**) Experimental setup for the study on insulin’s effect on gene expression patterns. Serum starved MCF7 cells were treated with DMSO (ctrl), insulin or insulin and torin1 followed by isolation of polysome-associated (P) and total (T) mRNA and their quantification using DNA-microarrays or RNAseq. (**B**) Principal component analysis using normalized DNA-microarray data as input (circles indicate clusters corresponding to RNA source and treatment [left]); and TMM-log2 RNAseq data (lines connect samples for each RNA source from the same replicate, i.e. illustrates a batch effect [right]). (**C**) Density plots of FDRs from anota2seq analysis of changes in polysome-associated mRNA, total mRNA, translational efficiency affecting protein levels and buffering using data from DNA-microarray or RNAseq data transformed with rlog or TMM-log2. (**D**) Density plots of mean expression levels under the control condition for mRNAs identified as changing their translation leading to altered protein levels using DNA-microarray and RNAseq data transformed with rlog or TMM-log2. Transcripts with FDR<0.05 that passed anota2seq filtering (material and methods) were included. Numbers of transcripts that overlap between platforms or data normalization/transformation (shared); or are unique are indicated.

As reported ^12^ principal component analysis of the DNA-microarray data set revealed the expected distinction between RNA source (polysome-associated or total) and treatment (**Fig. 4B**). The same analysis using RNAseq data revealed a similar separation based on RNA source but also a batch effect illustrated by a largely consistent pattern between the treatment groups within each replicate that was shifted across replicates (**Fig. 4B**). Such batch effects must be considered for efficient analysis ^52^. We therefore implemented a batch effect parameter in anota2seq, which is applied during APV and affects variance shrinkage. To assess the impact of the batch effect correction on anota2seq performance, we analyzed the RNAseq data with and without batch effect correction. We then selected mRNAs with an FDR < 0.05 (that also passed filtering criteria, see materials and methods) in analysis of changes in polysome-associated mRNA, total mRNA, translational efficiency affecting protein levels and buffering (anota2seq allows for differential expression analysis of polysome-associated or total mRNA using the same model as applied during analysis of translational efficiency affecting protein levels or buffering). When correcting for batch effects during anota2seq analysis, there is a substantial increase in the number of mRNAs with an FDR < 0.05 (**Fig. S9B**). Thus it is critical to consider batch correction during antoa2seq analysis ^52^. Notably, among the assessed methods only Xtail is not equivalent to and does not have the option to perform an analysis incorporating batch effects.

### RNAseq allows identification of changes in translational efficiencies altering protein levels across a broader dynamic range as compared to DNA-microarrays

We then characterized similarities and differences between anota2seq analysis of the DNA-microarray and RNAseq data. We first observed that RNAseq quantification (rlog or TMM-log2) leads to identification of more mRNAs showing differential polysome-association or total mRNA levels between control and insulin stimulated cells as compared to DNA-microarrays (**Fig. 4C**). We next collected anota2seq identified mRNAs that change their translational efficiency leading to changes in protein levels (FDR < 0.05 and passed filtering as indicated in materials and methods) when using DNA-microarray or RNAseq data as input. These sets were compared for their expression levels under the control condition to reveal differences in sensitivity that depend on the strength of the signal. RNAseq uniquely identified a set of mRNAs with low signals on the DNA-microarray. When comparing rlog to TMM-log2 RNAseq data, rlog identified a set of mRNAs with lower expression levels as compared to TMM-log2 (**Fig. 4D**).

In general, mRNAs that change their translational efficiency leading to altered protein levels are defined using both FDR and fold change (FC) thresholds ^53,54^. We therefore assessed the impact of such thresholds on anota2seq analyses of RNAseq or DNA-microarray data by comparing commonly or uniquely identified mRNAs across platforms and data normalizations/transformations using multiple thresholds (FDR<0.05; FC > 1.5; or FDR < 0.05 and FC > 1.5). TMM-log2 showed a larger overlap to DNA microarray as compared to rlog at FDR<0.05 although most mRNAs were distinct between TMM-log2 RNAseq and DNA-microarrays (**Fig. 5A**). The FC threshold dramatically reduced the overlap (**Fig. 5A**). To further illustrate this, volcano plots were generated showing mRNAs identified by anota2seq as changing their translation leading to altered protein levels (FDR < 0.05 and filtering) (**Fig. 5B**). Thus, although mRNAs selected solely by FDR in the TMM-log2 based and DNA-microarray anota2seq analysis show substantial overlap, fold-changes are not comparable across platforms and normalizations/transformations.

**Figure 5.**
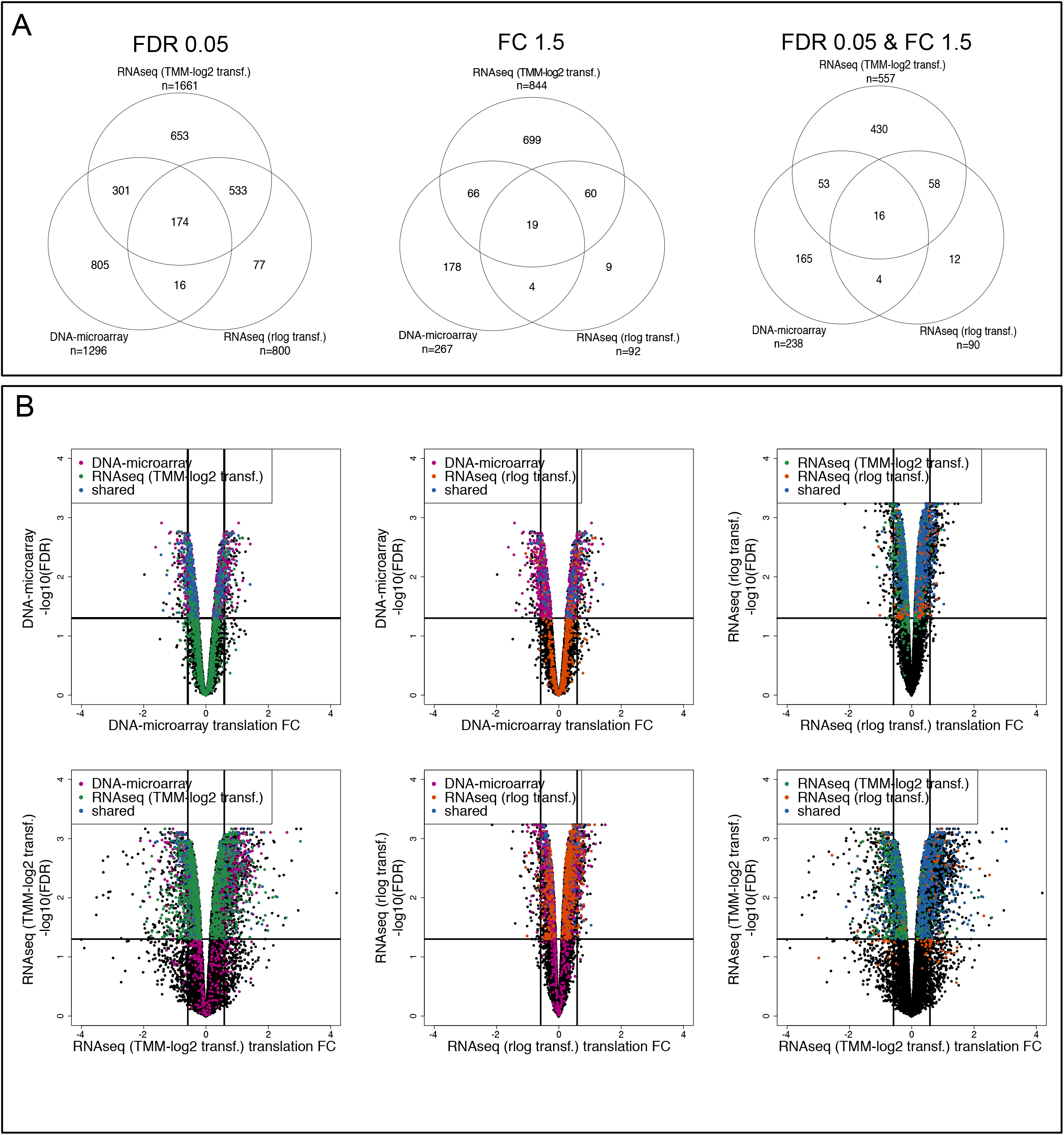
Fold-change filtering affects anota2seq analysis depending on platform and normalization/transformation. (**A**) Venn diagrams of mRNAs identified as changing their translational efficiency leading to altered protein levels by anota2seq using either FDR<0.05; FC>1.5; or FDR<0.05 & FC>1.5 thresholds and passing anota2seq filtering (see materials and methods). (**B**) Volcano plots of the anota2seq output for analysis of translational efficiency affecting protein levels using DNA-microarray data or RNAseq data from rlog or TMM-log2. Transcripts with FDR<0.05 that passed anota2seq filtering are included. Horizontal and vertical lines indicate 5% FDR or positive and negative log2 FC of 1.5 respectively. Unique and shared mRNAs are indicated as in figure 4D.

### Assessing the need for replication for efficient anota2seq analysis

As noted above, translatome data may include substantial variance; thereby highlighting that sufficient replication is essential for efficient analysis. Because this study includes four biological replicates and anota2seq requires two replicates when there are three treatment groups (3 replicates in the case of just two treatment groups), we determined the effect of reducing the number of replicates on anota2seq performance. Data sets were generated with all combinations of 2 and 3 replicates and analyzed using anota2seq to identify mRNAs changing their translational efficiency leading to altered protein levels (FDR<0.05 and filtering). When using 2 replicates per condition, we observed a drastic decrease in the number of identified mRNAs compared to scenarios where 3 or 4 replicates were used (**Fig. S9C**). We next monitored the FDRs of all mRNAs with an FDR < 0.05 (all replicates) during analysis using 2 or 3 replicates. This illustrated that with 3 replicates per condition, FDRs are below 0.05 for half of these mRNAs (**Fig. S9D**). Consequently, for efficient analysis using anota2seq, a minimum of 3 replicates should be applied - which is consistent with what has been proposed previously for analysis using babel ^29^. It should be noticed, however, that we used batch adjustment in the analysis, which reduces the degrees of freedom for the residual error in the APV ^24^ model. Fewer replicates may therefore be sufficient when analysing data sets without batch effects and/or with >3 conditions.

### Insulin modulates multiple steps in the gene expression pathway in a largely mTOR dependent manner

Next, we used anota2seq to determine the effects of insulin on gene expression in MCF7 cells. Gene expression was monitored at multiple levels including changes in mRNA abundance and translational efficiency leading to altered protein levels or buffering. This revealed that, for RNAseq based analyses, the major effect of 4 hour insulin treatment on gene expression occurs at the level of mRNA abundance followed by changes in translational efficiency altering protein levels (**Fig. 6A**). For DNA-microarray based analysis, as this analysis detected fewer changes in mRNA abundance, changes in translational efficiency leading to altered protein levels was more pronounced than changes in mRNA abundance (**Fig. 6A**). In addition to these modes of regulation of gene expression, we detected a sizable proportion of genes as translationally buffered (496 genes in the RNAseq TMM-log2 analysis). This suggests that translational buffering plays a major role in insulin-induced alterations in gene expression. Moreover, a comparison to the number of regulated mRNAs in insulin and torin1 vs vehicle treated cells revealed that most of the effects of insulin on gene expression are reversed by torin1 (**Fig. 6A**).

**Figure 6.**
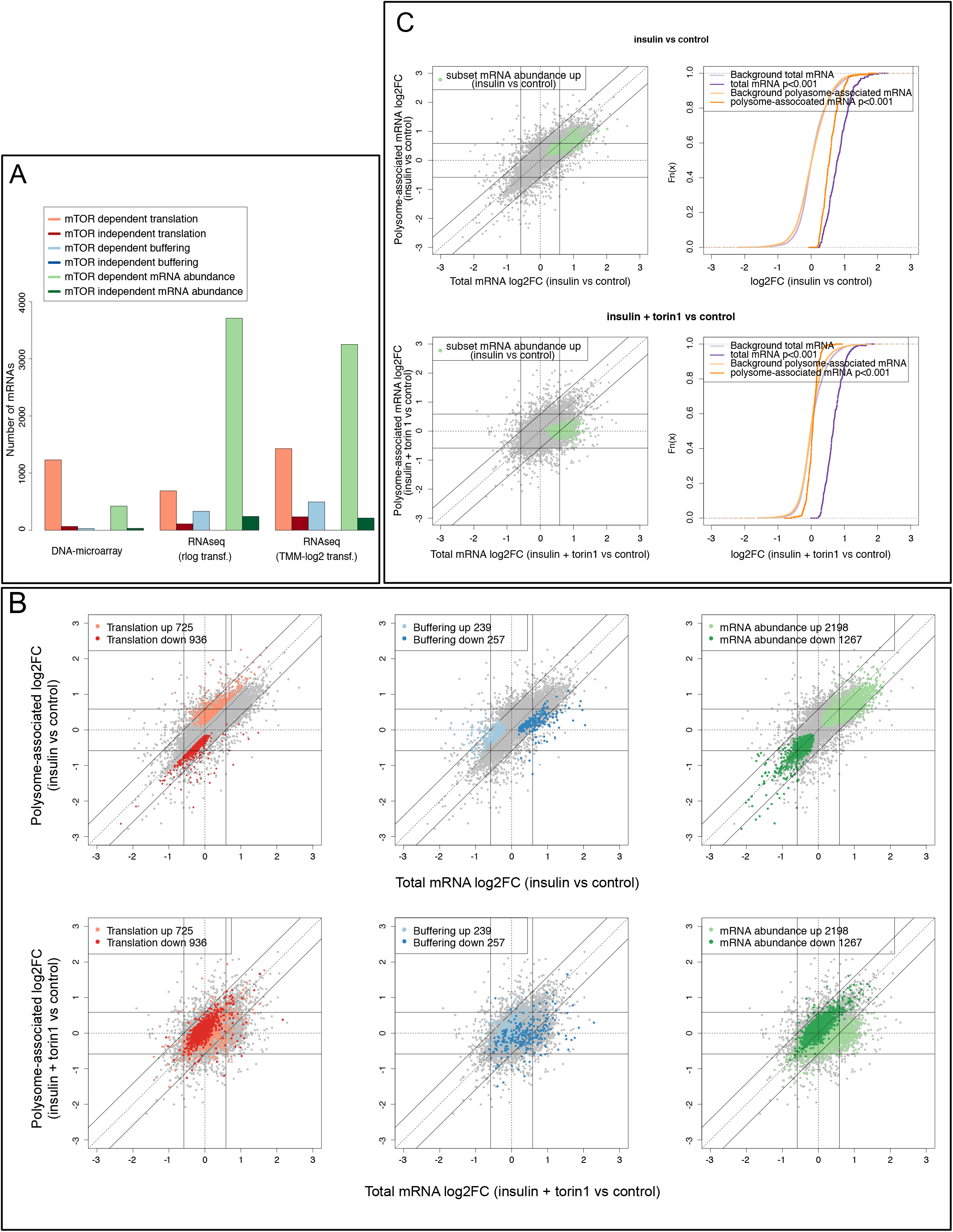
Insulin regulates gene expression at multiple levels in an mTOR dependent manner. (A) Numbers of mRNAs identified as changing their translational efficiency leading to altered protein levels or buffering, or mRNA abundance following insulin treatment as analyzed by anota2seq. The number in each category that did not depend on mTOR (i.e. not reversed by torin1) are also indicated. (B) Scatter plots of fold-changes for polysome-associated and total mRNA levels for the comparisons between insulin vs control (top row) and insulin plus torin1 vs control (bottom row). Genes that were identified as changing their abundance or translational efficiency leading to changes in protein levels or buffering upon insulin stimulation are indicated for both comparisons. (**C**) Similar to (B) but only for the subset of mRNAs whose abundance was congruently increased by insulin and buffered down upon torin1 treatment (left). Cumulative distribution functions of fold changes for polysome-associated mRNA and total mRNA for the subset (i.e. from plots to the right) for insulin vs control (top) and insulin plus torin1 vs control (bottom) as compared to the background (i.e. not in subset).

### Suppression of mTOR activity during insulin stimulation leads to translational buffering

To further establish the role of mTOR in mediating the effects of insulin on changes in gene expression, we employed scatter plots to visualize mRNAs with an FDR < 0.05 (insulin vs control) whose expression was altered via changes in mRNA abundance, translation affecting protein levels or buffering (**Fig. 6B**). This revealed an almost complete reversal of the changes in gene expression induced by insulin in the presence of torin1. However, a subset of mRNAs that changed their abundance upon insulin stimulation (i.e. congruently for total and polysome-associated mRNA) appeared to be translationally buffered in the presence of torin1 inasmuch as their polysome occupancy was not altered despite changes in mRNA levels as compared to the control (**Fig. 6B**, upper and lower right side plots). To further characterize this unexpected effect, we generated the overlap between mRNAs whose abundance was induced by insulin (as compared to control) that were also buffered down in the presence of torin1 (as compared to control; 237 mRNAs). Scatterplots and cumulative distributions illustrated that insulin induces congruent changes in mRNA abundance and polysome-associated mRNA for these genes (**Fig 6C**). When insulin was combined with torin1, however these genes were translationally buffered as compared to the control (**Fig 6C**). Collectively these data demonstrate that insulin induces total mRNA levels of a subset of mRNAs independently of mTOR but, in contrast, the translation of these mRNAs is mTOR-dependent. This leads to translational buffering of these mRNAs in insulin plus torin1 treatment vs. control conditions. As a consequence, although mRNA levels encoded by a subset of translationally buffered genes are higher in insulin plus torin1 treated relative to the control cells, it is expected that the levels of corresponding proteins will be comparable under both conditions. This suggests that for a subset of insulin induced genes, mTOR uncouples mRNA and protein levels via translational buffering.

### Insulin and mTOR coordinate expression of functionally related genes predominantly at the level of translation

We next sought to determine differences between quantification of polysome-profiling using DNA-microarray and RNAseq in enrichment of biological functions among identified genes showing insulin sensitive expression. Initially we compared the number of gene ontology (GO) terms identified for each platform and data normalization/transformation (FDR<0.01). This revealed that the majority of GO terms enriched among genes regulated using DNA-microarray data were also enriched using RNAseq data (**Fig. 7A**). Furthermore, RNAseq identified an additional set of GO terms as enriched (**Fig. 7A**). Identification of additional GO terms by RNAseq could be due to the larger number of identified genes (**Fig. 6A**). Alternatively, higher amounts of false positives may hamper GO analyses leading to identification of fewer GO terms under DNA-microarray based quantification. To distinguish between the two possibilities, we randomly sampled the same number of genes for each regulatory pattern (changes in translational efficiency altering protein levels or buffering; and mRNA abundance) and repeated the GO analysis (**Fig. 7B**). Indeed, application of a GO analysis on sets with equal amounts of genes results in identification of much fewer additional GO terms from analyses using RNAseq as input data. Thus, RNAseq allows identification of additional enriched biological processes by providing a larger set of regulated genes.

**Figure 7.**
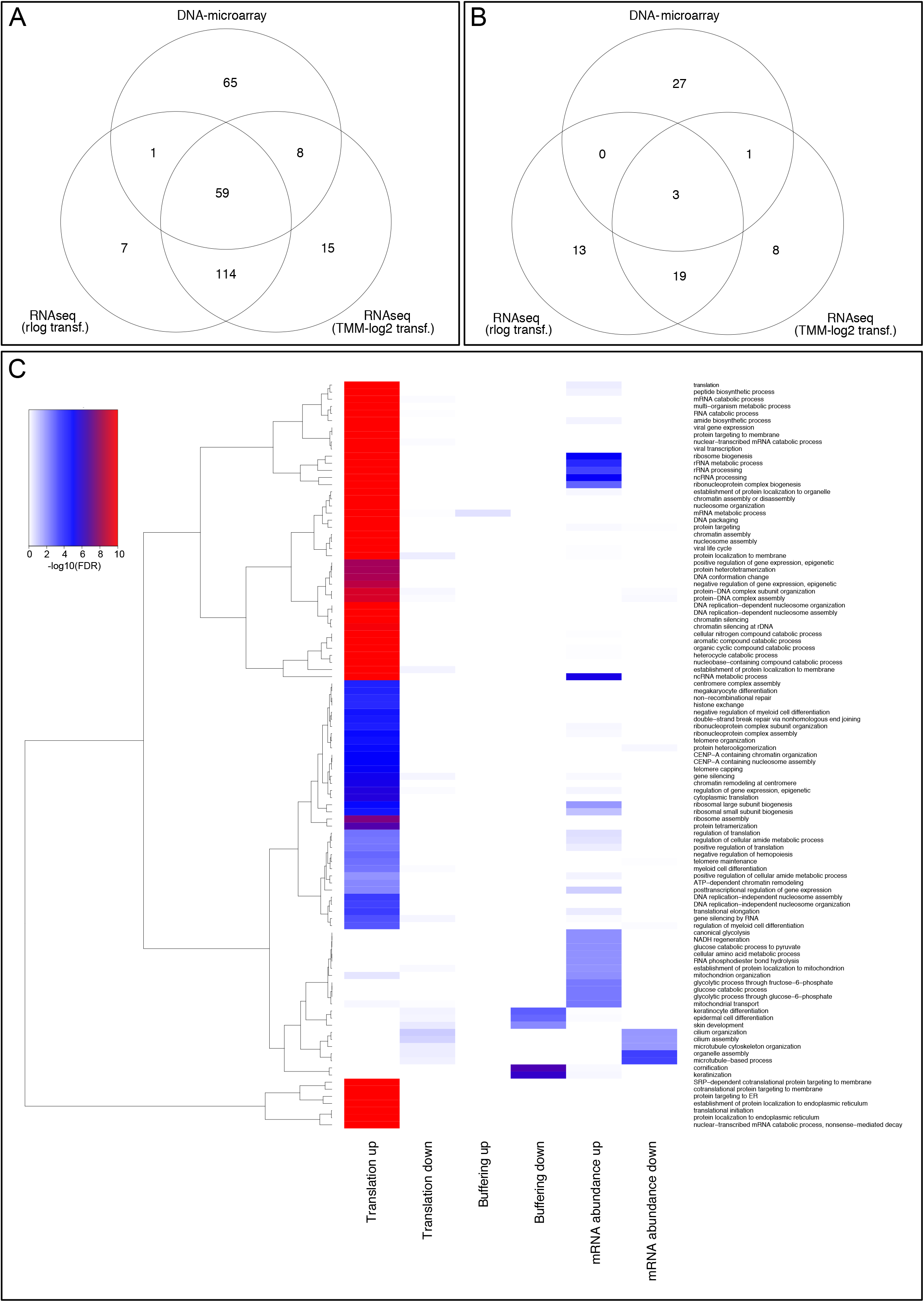
Changes in translational efficiency and mRNA abundance downstream of insulin signalling target distinct biological processes. (**A**) Venn diagram of enriched biological processes identified using all significant genes from all regulatory modes separately comparing insulin to control. (**B**) Same as (A) but with the same number of genes sampled to each regulatory mode prior to analysis (i.e. to have identical sizes of the regulated sets of genes). (**C**) Heatmap of –log10(FDR) values of significantly (FDR < 0.01) enriched biological processes identified using the TMM-log2 RNAseq data in the insulin to control comparison.

To explore whether different mechanisms regulating gene expression co-ordinate distinct functions we focused on the TMM-log2 RNAseq data (identified genes for each regulatory pattern can be found in **Supplemental File 5**). This was based on the finding that such data allows for detection of a higher number of regulated mRNAs (**Fig. 6A**) likely due to the larger dynamic range for fold-changes (**Fig. 5B**). We then compared overrepresented GO terms (FDR < 0.01) from sets of genes showing changes in translational efficiency altering protein levels, buffering and mRNA abundance (**Supplemental File 6**). The sets of mRNAs showing changes in translational efficiency affecting protein levels were enriched for a larger non-overlapping set of GO terms compared to those regulated via congruent changes in polysome-associated mRNA and total mRNA levels (**Fig. 7C**), despite that the latter subset contained a larger number of transcripts (**Fig. 6A**). This suggests that although our studies indicate more wide-spread regulation of total mRNA levels, translational efficiencies are modulated to coordinate expression of functionally related genes in a more focused manner. Thus, insulin regulates gene expression by modulating multiple mechanisms that target genes with distinct functions.

## DISCUSSION

The importance of mRNA translation in regulating gene expression has been exemplified in numerous biological and pathological processes ranging from stress response and cancer ^55^ to learning and memory ^56^. Identification of specific mRNAs that are under translational control requires interrogation of translatomes. Translatomes are vastly under-studied in comparison to transcriptomes (which reflect mRNA abundance determined at the level of transcription and/or mRNA stability) ^7^. Further research is thus required to understand the full complexity of translatomes and their impact on cell physiology and pathology. Such understanding largely relies on stringent and efficient application of transcriptome-wide methods to measure changes in translational efficiency. Moreover, downstream of methods applied to generate such data, sufficient replication and efficient data analysis is crucial for deriving valid conclusions. Notwithstanding that the noisy nature of polysome- and ribosome-profiling data is widely acknowledged, it is often paradoxically suggested that methods for identification of differential translation that do not require replication are warranted ^28^. This does not seem to be consistent with concerns about reproducibility in quantitative biology, which instead suggests that sufficient replication is essential to derive meaningful conclusions.

Currently, polysome- and ribosome-profiling are most commonly used to interrogate translatomes on a transcriptome-wide scale whereby polysome-profiling is more efficient for identification of changes in translational efficiency ^11,22^, while ribosome-profiling generates genome-wide information about ribosome positioning ^14,15^. A powerful unique property of polysome-profiling is that it allows examination of 5’ and/or 3’UTRs of mRNAs that are translated thereby facilitating identification of regulatory elements as well as potential differences in translation of mRNA isoforms that are co-expressed but differ in their 5’ or 3’UTRs ^11^. Thus, ribosome- and polysome-profiling represent compatible methodologies that provide ample opportunity to study translatomes. There is therefore a heightened interest to develop efficient methodologies to analyze polysome- and ribosome-profiling data. Such analysis need to be adapted to advances in technology that bear distinct characteristics, such as the count nature of RNAseq data, but also has to parallel the understanding of mechanisms regulating mRNA translation. Translational buffering represents one such emerging mechanism of translation control wherein alterations in mRNA levels are buffered at the level of translation such that levels of polysome-associated mRNAs are not affected by the changes in mRNA abundance ^17,37,41^. Such regulation retains protein levels but the mechanisms, extent and importance of this phenomenon are yet to be established. Nevertheless, translatome analysis requires efficient distinction between changes in translational efficiency that alter protein levels and buffering.

To address this major issue in studying translatomes, we developed the anota2seq algorithm, which can be employed to analyze DNA-microarray and RNAseq data and efficiently identify and separate changes in translational efficiency affecting protein levels from buffering. Evaluation of anota2seq compared to other methods for translatome analyzes indicated superior performance of anota2seq in detecting differential translation with low type-1-error rates and robustness against noise and varying sequencing depths. One unexpected finding from these studies was the poor performance of Xtail under the NULL condition inasmuch as a large number of mRNAs were identified as differentially translated despite no true changes in their translational efficiency (Fig. 1C and 2D). This possibly stems from the usage of the same data multiple times during Xtail analysis (e.g. both polysome-associated mRNA and the ratio between polysome-associated RNA and total mRNA) thereby not fulfilling the need for statistical independence. This underlines the importance of assessing the performance under the NULL condition during method development to derive tools that can be used for efficient and valid analysis. Moreover, anota2seq has several distinct features as compared to other methods: i) it is not based on interpretation of difference between log ratios and hence will not be affected by spurious correlations; ii) it distinguishes changes in translation efficiency affecting protein levels from buffering; iii) it allows for batch correction; and iv) it permits analysis of polysome-associated and total mRNA changes using the same analytical methods thereby allowing for simple and comparable identification of changes in polysome-associated mRNA, total mRNA, translational efficiency affecting protein levels and buffering. Our comparisons between data obtained from DNA-microarrays and RNAseq transformed by rlog or TMM-log2 suggest that TMM-log2 is preferred based on its larger dynamic range. Nevertheless, rlog and TMM-log2 are themselves comparable and a major impact on the overlaps is the estimated fold-changes where rlog is associated with smaller fold-changes as compared to DNA-microarrays and TMM-log2. Thus, due to technological biases and differences in data normalization/transformation methods, prudence is advised when selecting normalization/transformation and filtering thresholds for differential expression analysis. Anota2seq therefore incorporates both TMM-log2 and rlog but also allows the user to supply custom transformed and normalized data. Although we assessed the performance of anota2seq for analysis of polysome-profiling data, it is equally applicable to data from ribosome-profiling.

We ^57^ and others ^40,58^ have recently reported that mTOR primarily regulates gene expression by modulating mRNA translation. Our present study suggests that while insulin alters mRNA translational efficiencies affecting protein levels in an mTOR-dependent manner, mTOR also appears to mediate the majority of the effects from insulin on total mRNA levels. Moreover, we illuminate a previously unrecognized role of mTOR in regulating gene expression via translational buffering by showing that reduced mTOR activity buffers mTOR-independent changes in mRNA levels induced by insulin. The net outcome is likely unchanged protein levels despite alterations in mRNA levels. This suggests a fail-safe mode of regulation similar to what is commonly observed during immune responses where signalling activating both transcription and translation is needed to induce changes in the proteome ^7^. Furthermore, it appears that translational buffering is not limited to insulin stimulation inasmuch as a similar phenomenon was observed in response to LPS ^41^. This indicates that this mechanism plays a major role in perturbations in gene expression in response to a variety of stimuli. Moreover, this study advances the translational buffering concept by providing mTOR as the first pathway that can decouple polysome-associated and total-mRNA levels via this mechanism. These findings also unravel the complexity of regulatory networks, whereby insulin regulates gene expression at multiple levels. Understanding how regulation of total mRNA, mRNA translation and translational buffering is orchestrated, requires future studies where changes in gene expression will be monitored at different time points. In summary, we designed anota2seq for analysis of changes in translational efficiency affecting protein levels and buffering which can be applied independent of platform used for quantification. Application of such statistically stringent analyses holds a substantial promise to facilitate our understanding of dynamic regulation of translatomes in health and disease.

## MATERIALS AND METHODS

### RNA isolation for polysome-profiling

MCF7 cells were obtained from American Type Culture Collection (ATCC) and maintained in RPMI-1640 supplemented with 10% fetal bovine serum, 1% penicillin/streptomycin and 1% L-glutamine (all from Wisent Bio Products). Cell lines used were verified by profiling 17 short tandem repeat (STR) and 1 gender determining loci using ATCC Cell Line Authentication Service and the results were as following: MCF7 (100% match to ATCC MCF7 cells # HTB-22): D5S818 (11, 12 versus 11, 12); D13S317 (11 versus 11); D7S820 (8, 9 versus 8, 9); D16S539 (11, 12 versus 11, 12); vWA (14, 15 versus 14, 15); THO1 (6 versus 6); AMEL (X versus X); TPOX (9, 12 versus 9, 12), CSF1PO (10 versus 10). Cells were serum-starved for 16h followed by 4h treatment with either DMSO, insulin (4.2nM) or insulin (4.2nM) + torin1 (250nM; cells were pre-treated for 15 minutes with torin1 before stimulation with insulin), washed with ice cold PBS containing 100µg/mL cycloheximide, collected and lysed in a hypotonic lysis buffer [5 mM Tris-HCl (pH 7.5), 2.5mM MgCl2, 1.5 mM KCl, 100 µg/mL cycloheximide, 2mM DTT, 0.5% Triton X-100, and 0.5% sodium deoxycholate] (all chemicals used were obtained by Sigma unless otherwise stated). A lysate sample was collected and total RNA was isolated using TRIzol (Invitrogen). Lysates were loaded onto 10–50% (wt/vol) sucrose density gradients [20 mM Hepes-KOH (pH 7.6), 100 mM KCl, 5 mM MgCl2] and centrifuged at 36,000 rpm [SW 40 Ti rotor (Beckman Coulter, inc)] for 2h at 4 °C. Gradients were fractioned and the optical density at 254 nm was continuously recorded using an ISCO fractionator (Teledyne ISCO). RNA from each fraction was isolated using TRIzol (Invitrogen). Fractions with mRNAs associated with >3 ribosomes were pooled (polysome-associated mRNA). The experiment was performed in 4 biological replicates.

### Protein isolation and Western Blotting

MCF7 cells were serum-starved overnight with RPMI supplemented with 0.1%FBS. Where indicated, cells were pre-treated with torin1 (250nM) for 15 min and stimulated for 4 hours with insulin (4.2nM). Cells were washed twice in ice-cold PBS and lysed on ice in RIPA buffer [20 mM Tris-HCl (pH 7.5), 150 mM NaCl, 1 mM Na2EDTA, 1 mM EGTA, 1% (v/v) NP-40, 1% (w/v) Na-deoxycholate, 0.1% (w/v) SDS, 2.5 mM Na-pyrophosphate, 50 mM NaF, 20 mM β-glycerophosphate, 1mM Na3VO4] supplemented with 1X complete protease inhibitors (Roche). Protein concentrations were determined using Pierce BCA Protein Assay Kit (Thermo Fisher Scientific) and lysates were prepared in 4X Laemmli buffer [250mM Tris/SDS (pH 6.8), 40% glycerol, 8% SDS, 400mM DTT and 0.01% bromophenol blue]. Proteins of interest were detected using the following primary antibodies: anti-β-actin (Clone AC-15) #A1978 from Sigma used at 1:5,000 dilution; and anti-4E-BP1(53H11) #9644, anti-p-4E-BP1 (S65) (174A9) #9456, anti-rpS6 #2217, anti-p-rpS6 (S240/244) #2215, anti-S6K1/2 #9202, anti-p-S6K1/2 (T389) #9205 all used at 1:1,000 dilutions from Cell Signaling Technologies. Secondary antibodies (Amersham) were used at 1:10,000, and signals were revealed by chemiluminescence (ECL, GE Healthcare) on X-ray blue film.

### DNA-microarray data processing

DNA-microarray based expression analysis (polysome-associated and total RNA) was performed using the Human Gene ST 1.1 array plate (Gene Titan from Affymetrix) by the Bionformatics and Expression Analysis (BEA, Karolinska Institutet, Stockholm). Gene expression was summarized and normalized using Robust Multi-array Average (RMA) ^59^ using custom probe set annotation (hugene11st_Hs_ENTREZG) as these provide improved precision and accuracy ^60–62^. Microarray quality control was performed using standard methods ^63^ and all arrays were considered of high quality. Raw data is available at the Gene Expression Omnibus (GSE76766).

### RNAseq data processing

RNAseq libraries were obtained according to the strand specific TruSeq protocol (Illumina) including ribosomal RNA depletion (RiboZero) and sequenced using the Illumina HiSeq 2000 platform obtaining single end 50 nucleotide reads for both polysome-associated and total RNA. Library preparation and sequencing were performed at the Science for Life Laboratory Genomics Facility (Karolinska Institutet, Stockholm). RNAseq reads were subjected to and passed quality control (Phred scores >30) and were subsequently aligned to the hg19 reference genome using HISAT ^64^ with default settings. The average percentage of reads mapped to the genome was 83% (average number of reads was 92 million). Reads mapped to multiple locations in the genome were discarded. Gene expression was quantified using the RPKMforgenes.py method ^65^ from which raw gene counts were obtained (version last modified 11.04.2013). Genes that could not be resolved (based on sequence similarity) and/or had 0 counts in at least one sample were removed from the analysis. Genes were annotated using the RefSeq ^66^ database. A total of 12,252 unique genes were represented in the RNAseq data. Raw counts were either rlog ^30^ transformed using default settings; or normalized using TMM normalization ^39^ and log2 transformed using the voom R-package ^67^ with default settings (such data is referred to as TMM-log2 below).

### RNAseq data simulation using empirical data

Method performance comparisons were done using simulated data. Data was simulated by sampling from a negative binomial (NB) distribution using estimated means and dispersions from empirical RNAseq data produced in this study (control and insulin conditions). We used a similar approach as described previously ^68^. For polysome profiling data, in addition to unchanged mRNAs, three regulated groups were simulated: translation affecting protein levels (a change in polysome-associated mRNA independent of changes in total mRNA; **Fig. 1A**); buffering (a change in total mRNA that is not reflected by a similar change in polysome-associated mRNA; **Fig. 1A**); and mRNA abundance (i.e. a congruent change in polysome-associated and total mRNA; **Fig. 1A**). Each regulatory group is represented by 5% of all mRNAs (total number of simulated mRNAs was 10,748). Four replicates of control and treatment conditions were simulated using the following parameterization for the NB distribution:

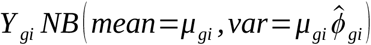

where *Y_gi_* is the simulated count for gene *g* and RNA source *i* (i.e. polysome-associated or total mRNA) *and ϕ*_*gi*_ is the dispersion. The mean *μ_gi_* and dispersion *ϕ_gi_* were estimated from the empirical data using maximum likelihood estimates 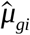 and 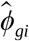 with

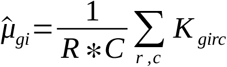

where R is the number of replicates, C the number of conditions and K*_girc_* the read count for the empirical gene *g*, RNA source *i*, replicate *r* and condition *c*. 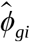 was obtained by maximizing the log-likelihood function described ^68^. 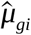 and 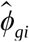 were then directly used as parameters for a NB distribution to simulate 4 replicates of the control condition. We used the rnbinom function of the stats R package with parameter size of 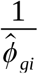.

For unchanged mRNAs (i.e. not belonging to any of the regulated groups), the treatment condition was sampled from a distribution with 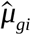 and 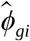 identical to the control condition for that gene. For regulated mRNAs, these estimates were used as base parameters and then modified: For the translation affecting protein levels group (**Fig. 1A**; 537 mRNAs, similar number of up- and down-regulation), the base parameters were used to simulate total mRNA for both conditions and polysome-associated mRNA for the control condition. The mean and dispersion parameters used to simulate polysome-associated mRNA under the treatment condition were modified as follows: an effect parameter *α_g_* for upregulation was sampled from a vector containing values from 1.5 to 3 with steps of 0.2. For down regulation the effect parameter was modified to 1/ *α_g_*. The modified mean parameter was then 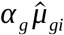 (or 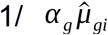 for down regulation). In order to keep the mean-variance relationship as similar as possible to the empirical data, the modified dispersion was taken as the dispersion of the gene from the empirical data having the closest mean estimate to 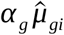 (or 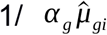 for down regulation). Similarly, for the buffering group (**Fig. 1A**; 537 mRNAs), base parameters were used for polysome-associated mRNA (both conditions) and total mRNA (control condition). A random effect was introduced for simulating total mRNA under the treatment condition and applied as described for the translation group. Finally, for the mRNA abundance group base parameters were used for total and polysome-associated mRNA under the control condition and the same effect was introduced to modify the parameters of the distribution from which total and polysome-associated mRNA were sampled under the treatment condition; and applied as described for translation. Transcripts that had a simulated count of zero in any sample were removed before analysis (To assess the variability of the simulation, 5 such data set were simulated and an example sampled data set is supplied as **Supplement File 1**).

To assess the ability of methods to control for type I error/false discovery rate in the absence of any true regulation between control and treatment, we also simulated a null dataset using base parameters for all mRNAs in both conditions (i.e. all mRNAs are unchanged; the NULL data set is supplied as **Supplement File 2**). For further evaluation, we simulated datasets with additional 5%, 10% and 15% noise (where the noise is a percentage of each count, which is added or subtracted [same probability to add or subtract]) and datasets with different sequencing depths (5, 10, 15 and 25 million RNAseq read counts per sample).

### Comparison of methods for analysis of differential translation using simulated data

We compared anota2seq to four tools available in the statistical programming language “R” for analysis of the translatome: babel ^29^ (version 0.3.0), DESeq2 ^30^ (version 1.10.1), translational efficiency score (TE score) and Xtail ^28^ (version 1.1.3). All analyses were performed using R (version 3.2.3). Simulated raw count data were used as input for babel ^29^, DESeq2 ^30^ and Xtail ^28^. For TE score analysis, counts were normalized using DESeq2 (normalization for library size using the median ratio method) ^69^ and log2 transformed. For anota2seq, counts were either rlog ^30^ or TMM-log2 ^39^ transformed. Similar to anota ^23^, anota2seq combines analysis of partial variance (APV) ^24^ and the Random Variance Model (RVM) ^25^ and uses a two-step process that firstly assesses the model assumptions for 1) absence of highly influential data points, 2) common slopes of sample classes, 3) homoscedasticity of residuals and 4) normal distribution of per gene residuals. This is followed by analysis of changes in translational efficiency affecting protein levels or buffering using APV ^24^ and RVM ^25^. Babel ^29^ uses an errors-in-variables regression consisting of two steps. Using a NB distribution in both steps, total mRNA levels are modelled as a first step followed by modelling of polysome-associated mRNA as a second step. To estimate the mean in the second part of the model a trimmed least-squares approach is used. Following a parametric bootstrap, for every gene, a test is performed for the null hypothesis that observed polysome-associated mRNA levels are as expected from total mRNA levels. Per gene per sample p-values are then combined into one single p-value using the arithmetic mean of p-values. DESeq2 ^30^ applies a generalised linear model approach. For all analyses done in this study we used the following model:

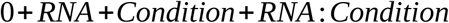

Here RNA is either polysome-associated or total mRNA; condition is control or treatment; and RNA:Condition the interaction effect. In this model RNA:Condition is the measurement for differential translation. The TE score is the difference between conditions in log2 ratios (between polysome-associated and total mRNA). Statistics for changes in TE were calculated using Student’s t-test (p-values were adjusted using the Benjamin-Hochberg approach) ^70^. Xtail ^28^ is a method based on comparisons of fold-changes between conditions using data from polysome-associated mRNA, total mRNA and ratios between polysome-associated and total mRNA. For each comparison, a fold change probability distribution is generated. Using these probability distributions a joint probability matrix is derived and p-values for each gene assessed. For all methods, we used either the reported false discovery rates (FDR) for all mRNAs when applying default parameters or the selection of filtered mRNAs (filtered using default parameters for each method) under a specific FDR threshold. These tools were compared using receiver operator curves on simulated data. The outputs from all methods for the two different data sets described above are provided as **Supplemental Files 3–4**.

### Analysis of the empirical data set using anota2seq

DNA-microarray and RNAseq data were analyzed for changes in polysome-associated mRNA, total mRNA, translation affecting protein levels and buffering using the anota2seq algorithm. During the analyses of the RNAseq data, if not stated otherwise, the replicate was included in the model to account for a batch effect (as indicated in the results section). Transcripts where considered changing their translational efficiency leading to altered protein levels or buffering when passing filtering criteria in anota2seq (minSlope = −0.5; maxSlope = 1.5, deltaPT = 0.15 (for translation) and minSlope = −1.5; maxSlope = 0.5, deltaTP = 0.15 (for buffering)) and significance criteria (three different were evaluated: maxRvmPAdj = 0.05 [i.e. FDR < 0.05]; minEff = log2(1.5) [i.e. abs(Fold Change) < 1.5]; maxRvmPAdj = 0.05 & minEff = log2(1.5)). These filtering criteria were used for both DNA-microarray and RNAseq analysis. All analysis and comparison of DNA-microarray and RNAseq data were done on mRNAs quantified on both platforms leading to a total of 10,957 mRNAs.

### R-packages and settings

Hierarchical clustering, scatter plots and PCA plots were generated using the hclust (default settings), smoothScatter and prcomp (center=TRUE) R-functions, respectively. Receiver operator curves (ROC), area under the curve (AUC), partial AUC (pAUC) and precision recall curves were generated using the ROCR R-package (version 1.0–7)^42^. P-values from each method were used in the ROC analysis. pAUC were obtained at 5 and 15% false positive rate (fpr) by using the fpr.stop parameter from the ROCR package; the so acquired AUC was divided by the corresponding fpr cut-off rate. The precision is the positive predictive value and the recall is the true positive rate or sensitivity. Biological process (BP) Gene Ontology (GO) analysis was performed using a hyper-geometric test for GO term overrepresentation using the GOstats R-package (2.42.0). Each set of regulated genes (changes in mRNA abundance or translational efficiency altering protein levels or buffering) separated based on directionality of regulation (i.e. up or down-regulated by insulin) was processed. The background was the set of quantified genes that overlapped between RNAseq and DNA-microarrays. Only GO terms with >10 and < 500 members out of which at least 10 were included in the set of regulated genes were considered. P-values of the filtered GO terms were then adjusted for multiple testing using the Benjamin-Hochberg approach ^70^.

### Anota2seq software

The anota2seq software is available as a Bioconductor package.

### Statistics

All statistical tests within the anota2seq package are 2-tailed.

## Acknowledgements

We acknowledge support from Science for Life Laboratory, the Knut and Alice Wallenberg Foundation, the National Genomics Infrastructure funded by the Swedish Research Council, and Uppsala Multidisciplinary Center for Advanced Computational Science for assistance with massively parallel sequencing and access to the UPPMAX computational infrastructure. This research was supported by the Swedish Research Council, the Swedish Cancer Society, the Cancer Society in Stockholm, the Wallenberg Academy Fellows Program, and STRATCAN grants (to O.L.); and Canadian Institutes for Health Research (PJT-148603) and Canada Cancer Society Research Institute Innovation Grant (703816) to IT. I.T. is a Junior 2 Research Scholars of the Fonds de Recherche du Québec – Santé (FRQ-S). MC is a recipient of Canadian Institutes for Health Research postdoctoral fellowship. LF is supported by the Department of Health and Human Services acting through the Victorian Cancer Agency (MCRF16007). IT acknowledges Prof. R. McInnes for invaluable advise and support.

## Author contribution

C. O., L. F., I. T. and O. L. designed the study. C. O., J. L., V. G., C. M., L. M., and M. C. performed experiments. C. O., J. L. and O. L. developed the anota2seq software. C. O., J. L., L. F., I. T., and O. L. analyzed and interpreted the data. C. O., J. L., I. T. and O. L. wrote the manuscript.

